# Latrophilins are essential for endothelial junctional fluid shear stress mechanotransduction

**DOI:** 10.1101/2020.02.03.932822

**Authors:** Keiichiro Tanaka, Andrew Prendergast, Jared Hintzen, Abhishek Kumar, Minhwan Chung, Anthony Koleske, Jason Crawford, Stefania Nicoli, Martin A. Schwartz

## Abstract

Endothelial cell (EC) responses to fluid shear stress (FSS) are crucial for vascular development, adult physiology and disease. PECAM1 is an important transducer but earlier events remain poorly understood. We therefore investigated heterotrimeric G proteins in FSS sensing. Knockdown (KD) in ECs of single Gα proteins had little effect but combined depletion of Gαi and Gαq/11 blocked all known PECAM1-dependent responses. Re-expression of Gαi2 and Gαq but not Gαi1 and Gαi3 rescued these effects. Sequence alignment and mutational studies identified that K307 in Gαi2 and Gq/11 (Q306 in Gαi1/3), determines participation in flow signaling. We developed pull-down assays for measuring Gα activation and found that this residue, localized to the GPCR interface, determines activation by FSS. We developed a protocol for affinity purification of GPCRs on activated Gα’s, which identified latrophilins (ADGRLs) as specific upstream interactors for Gαi2 and Gq/11. Depletion of latrophilin-2 blocked EC activation of Gαi2 and Gαq, downstream events *in vitro*, and flow-dependent vascular morphogenesis in zebrafish embryos. Surprisingly, latrophilin-2 depletion also blocked flow activation of two additional pathways activated at cell-cell junctions, Smad1/5 and Notch1, independently of Gα proteins. Latrophilins are thus central mediators of junctional shear stress mechanotransduction via Gα protein-dependent and -independent mechanisms.

## Introduction

Fluid shear stress acting on vascular endothelial cells (ECs) is a major determinant of vascular development, remodeling and disease^1^. ECs align in the direction of laminar shear stress via a pathway that involves cell-cell junctional proteins, integrins and Rho family GTPases ^2^. Regions of arteries where flow is disturbed, with lower magnitude and changes of direction during the cardiac cycle are susceptible to atherosclerosis, whereas regions of high laminar shear are highly protected^3^. While many pathways have now been catalogued that are regulated by fluid shear stress in ECs and progress has been made into their relevance to remodeling and disease, the fundamental molecular mechanisms that initiate signaling remain poorly understood.

We previously reported that the shear stress mechanotransduction is mediated by a complex of proteins at cell-cell contacts comprised of PECAM-1 (hereafter termed PECAM), VE-Cadherin (VEcad) and VEGF receptors (VEGFRs)^2,4^. PECAM is the direct mechanotransducer; thus, force application to PECAM triggers a subset of events activated by flow^5^, while its deletion blocks activation of flow-induced endothelial responses^2^. Application of force to PECAM in response to flow appears to occur not through direct force transmission, but rather by association of PECAM with the cytoskeleton, leading to force application from actomyosin^5^. These findings point toward an upstream signaling pathway that operates within seconds. GPCR/G protein signaling is an obvious candidate. Indeed, connections between GPCRs and shear stress have been reported^6-11^. However, the literature on this topic is highly inconsistent, with different GPCRs and mechanisms described in different papers.

Here we took an unbiased approach starting from the G*α* proteins. These studies led us to identify the adhesion type GPCR, latrophilins, as central upstream mediators of three unrelated flow sensing pathways operating at cell-cell junctions, mediated through both G protein-dependent and -independent mechanisms.

## Results

### Identification of G protein α subunits required for flow sensing

To investigate the roles of GPCR signaling in endothelial flow responses, we investigated the contributions of all of the main classes of G*α* proteins, Gs, Gi and Gq/11 and G12/13 using siRNA sequences designed to knock down (KD) all isoforms of each class. Single KD of each class had no or weak effects on EC alignment in flow (Gi and Gq/11: Figure 1A; Gs and G12/13: Extended data Fig1a&1b). However, combined KD of Gi plus Gq/11 completely blocked alignment (Fig 1A), whereas other combinations did not (Extended data Fig1c-e). PECAM and its downstream effectors are essential for EC alignment in flow, with src family kinases (SFKs) as the earliest known downstream signal^2^. Activation of VEGFRs and Akt are also downstream of PECAM. Single Gα KDs had little effect on flow activation of these events were strongly inhibited by KD of Gi plus Gq/11 (Figure 1B&C and Extended data Fig1f). Thus, G protein signaling is required for activation of multiple PECAM-dependent pathways.

**Figure 1.**
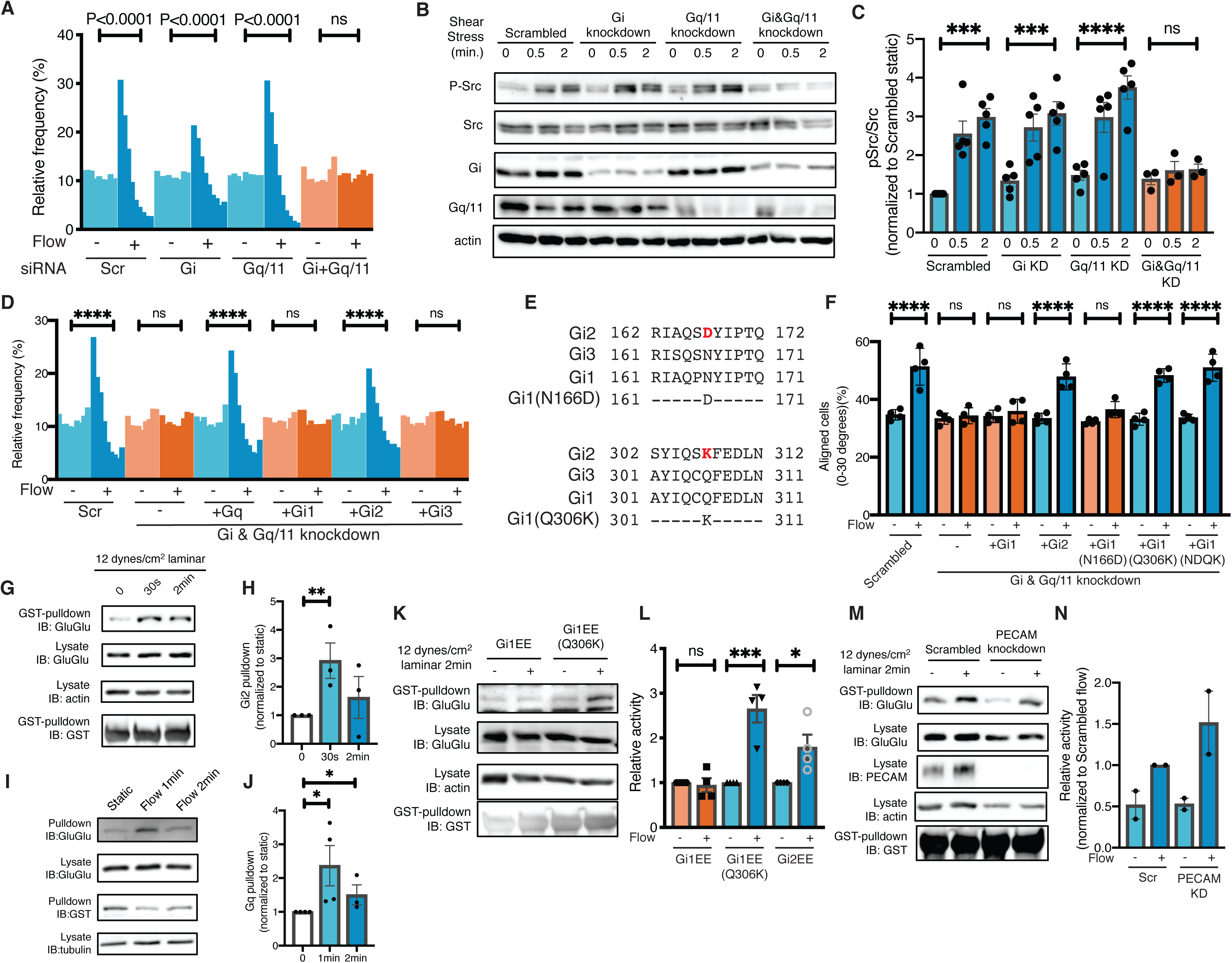
G proteins in endothelial flow responses. (**A**) HUVECs with KD of the indicated proteins were subjected to fluid shear stress (FSS) at 12 dynes/cm2 for 16 hours and nuclear orientation determined. Results are displayed as histograms showing the percent of cells within 10° of the direction of flow as described in Methods. Statistical analysis used one-way ANOVA with Tukey’s multiple comparisons test. (**B**) HUVECs after KD of the indicated proteins were analyzed by Western blotting for Src activation. (C) quantification of (B). values are means ± SEM. (**D**) Rescue of Gq/11/i KD by re-expression of siRNA-resistant versions of the indicated proteins. ****: p<0.0001, one-way ANOVA with Tukey’s multiple comparison test. (**E**) Amino acid sequences of Gi1, Gi2, and Gi3, and the Gi1 gain-of-function mutant. (**F**) Rescue of Gq/11/i KD with wild type Gi1, Gi2 and Gi1 gain-of-function mutants. Each point corresponds to one measurement averaged from >500 cells. N=4. ****: p<0.0001, one-way ANOVA with Tukey’s multiple comparisons test. (**G**) GINIP pulldown assay for activation of Gi2 by FSS. N=3. Results quantified in (**H**). **: p=0.0185, one-tailed Student’s t-test. (**I**) GRK2N pulldown assay for activation of Gq. N=4, quantified in (**J**). *: p<0.05, one-tailed Student’s t-test. (**K**) GINIP pulldown assay for activation of WT and Q307K Gi1. N = 4, quantified in (**L**). *: p=0.0304, ***: p=0.0017, two-tailed Student’s t-test. (**M&N**) Gi2 activation after PECAM KD. HUVECs expressing Gi2 GluGlu tag were transfected with scrambled siRNA or PECAM siRNA, then were exposed to FSS and Gi2 activation assayed as above, quantified in (**N**). Values are means ± range, N=2.

To address possible off-target effects, we re-expressed siRNA-resistant G*α* protein in KD HUVECs. As expected, siRNA-resistant Gq rescued cell alignment (Figure 1C and Extended data Fig 3a), however, among the Gi isoforms, we unexpectedly found that Gi2 but not Gi1 or Gi3 rescued (Figure 1D and Extended data Fig 3b). This result prompted us to examine the sequences of these α subunits for residues common to Gq, G11 and Gi1 but differing in Gi1 and Gi3. Two residues (using human Gi2 numbering throughout) fit this pattern: aa167 that is aspartate(D) in Gi2/q/11 but asparagine(N) in Gi1/3, and 307 that is lysine(K) in Gi2/q/11 but glutamine(Q) in Gi1/3 (Figure 1E and Extended data Fig 2). Rescue experiments showed that Q307K but not N167D Gi1 rescued cell alignment in flow (Figure 1F and Extended data Fig 3c). This residue therefore distinguishes the Gα subunits that do and don’t participate in FSS sensing.

### Novel G*α* activity assay shows flow-induced activation of heterotrimeric G*α* proteins

Structural and biochemical studies showed that K307 lies in the binding interface with GPCRs^12,13^, suggesting that it may determine which G*α* proteins are activated by flow. Testing this hypothesis therefore requires measuring G*α* activation in live cells. Despite the widespread use of indirect assays such as cAMP or calcium, direct whole cell assays are limited. We therefore developed pull-down assays based on specific binding of GTP-loaded Gα subunits to effector proteins (Extended data Fig 4a). The effectors GINIP and the GRK2 N-terminal domain were used for Gi and Gq, respectively^14,15^. To facilitate these assays and distinguihs Gα isoforms without specific antibodies, we prepared versions containing an internal GluGlu (EE) tag that does not perturb function (Extended data Fig 4b)^16,17^. Control experiments used constitutively active forms of G*α* proteins or the artificial DREADD GPCR that activates these Gα proteins. For Gq, early experiments showed weak responses (not shown) but co-expression of Ric8A, which stabilizes active Gq^18^, greatly amplified the signals. We found that active but not inactive Gi and Gq bound these effector proteins and that established ligands triggered activation (Extended data fig 4c-f). These novel assays revealed rapid activation of Gi2 and Gq by FSS (Figure 1G-J). Flow failed to activate Gi1 but Gi1Q306K was activated (Figure 1K & 1L). Gi2 activation by FSS was unaffected by PECAM KD, indicating that this step was upstream or independent of PECAM-1 (Figure 1M & 1N). Thus, flow activation of specific Gα subunits matches their requirement in flow sensing, both determined by residue 307.

### Identification of the upstream GPCR

We next sought to exploit this information to identify the GPCR(s) that mediates Gi2 and Gq/11 activation by flow. However, G*α* proteins dissociate from GPCRs immediately after GTP loading, making affinity purification problematic. We therefore made WT and Q307K Gi1 with the 4 alanine insertion (hereafter ins4A) in helix *α*5 that increases affinity for GPCRs even in the presence of GTP^19^. These G*α*(ins4A) mutants, which also contained an inserted GFP to facilitate affinity chromatography, localized predominantly to plasma membranes as do WT G proteins (Extended data fig 5a). Detergent inactivates GPCRs, thus, could be used in these assays. Instead, we solubilized with a recently developed non-detergent, nanodisc-forming reagent, styrene maleimide anhydride (SMA)^20^ (Figure 2A). To test this approach, we co-expressed these G*α* proteins with DREADD GPCR, extracted membranes with SMA, and isolated G*α* proteins using GFP-TRAP™ nanobody beads. The G*α* subunits co-purified with DREADD following receptor activation, validating this method (Extended data fig 5b&5c). Next, HUVECs expressing WT or Q306K Gi1(ins4A)-GFP were subjected to FSS, extracted with SMA and purified on GFP-TRAP™ beads (Extended data fig 5d). Proteomic analysis identified two GPCRs, S1PR1 and ADGRL3, that were specific to the Q306K mutant (Extended data fig 5e). S1PR1, however, is known to activate Gi1 and 3 as well as 2^21^, thus, has the wrong specificity. We therefore focused on ADGRLs, also called latrophilins (LPHNs).

**Figure 2.**
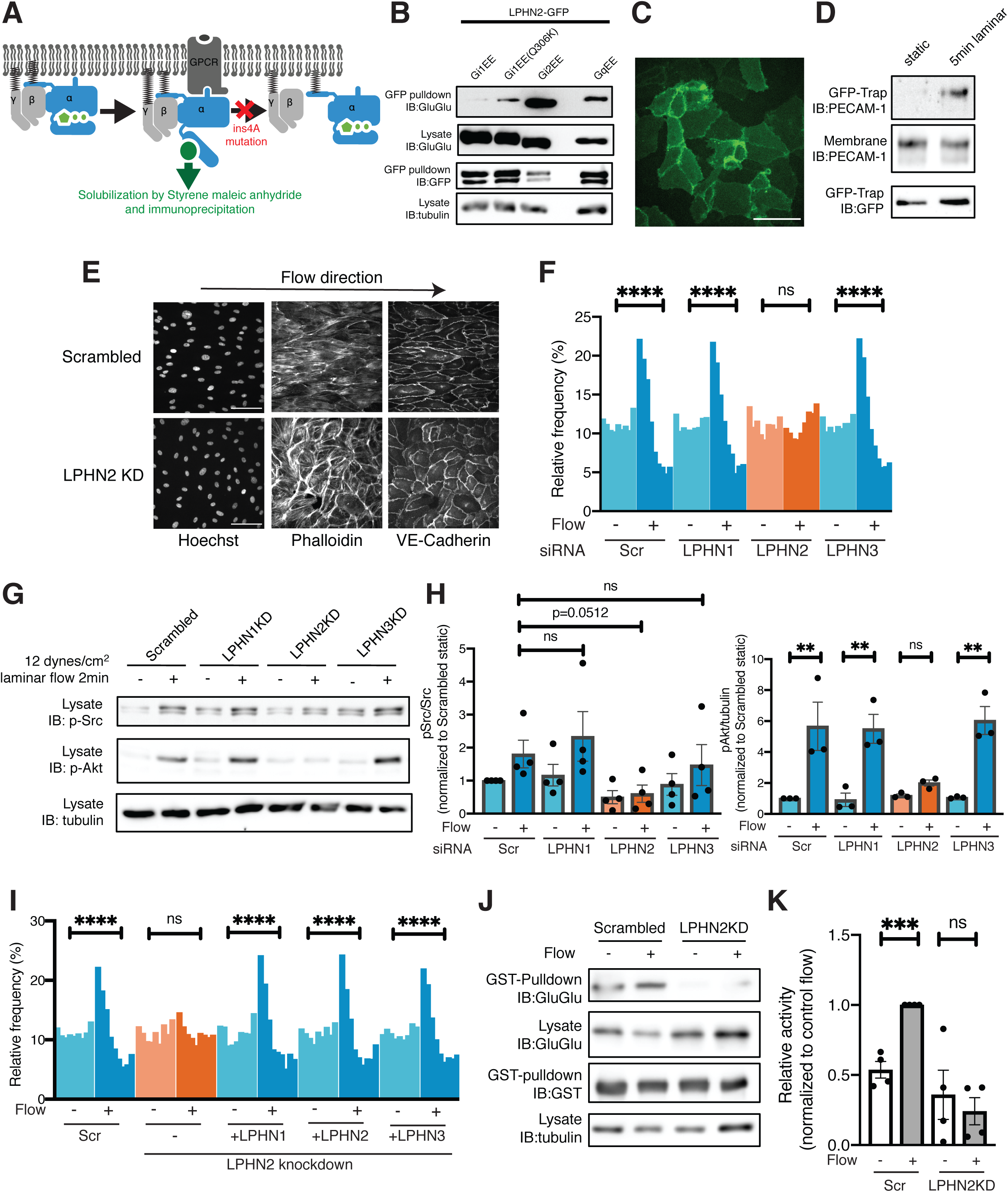
Identification of the upstream GPCR. (**A**) Strategy for proteomics with GPCR-trapping mutant of WT vs K306K Gi1. (**B**) Co-immunoprecipitation of LPHN2-GFP with indicated G proteins containing internal GluGlu tags. (**C**) Subcellular localization of LPHN2-mClover3. Scale bar: 100*µ*m. N=6. (**D**) Gi1(Q306K) pull-downs from ECs with and without 5 min FSS were probed for PECAM. (**E**) HUVECs after KD of individual latrophilin isoforms were subjected to FSS for 16 h, then fixed and stained with Hoechst, Phalloidin and antibody to VE-Cadherin. Scale bar: 100*µ*m. (**F**) Cell alignment quantified as in Fig 1 from >2000 cells/experiment, N=3. ****: p<0.0001, one-way ANOVA with Tukey’s multiple comparisons test. (**G**) Control and LPHN2 KD HUVECs assayed for activation of Src family kinases and Akt after FSS. N=3-4, quantified in (**H**). (**I**) LPHN2 KD HUVECs were rescued by re-expression of the indicated LPHN isoforms. ****: p<0.0001, one-way ANOVA with Tukey’s multiple comparisons test. Values are averaged from >2000 cells/experiment, n=2. (**J**) Gi2 pulldown assay after KD of LPHN2. N= 4. Results quantified in (**K**). ***: p<0.001, two-tailed Student’s t-test. Quantified values are means ± SEM throughout.

Latrophilins (1,2 and 3 in mammals) are adhesion-type GPCRs that regulate neuronal synapses^22^. LPHN1 and LPHN3 are found mainly in neurons, whereas LPHN2 is more widely expressed and is the major isoform in HUVECs based on the Gene Expression Omnibus (accession no. GSE71164). To confirm proteomic results, we immunoprecipitated the EE-tagged G*α* proteins (Figure 2B). Western blotting demonstrated that Gi2, Gq, and Gi1Q306K but not WT Gi1 pulled down LPHN2. GFP-tagged LPHN2 localized to cell-cell contacts, consistent with its connection to PECAM-1 (Figure 2C). Interestingly, proteomic analysis also detected PECAM-1 in the Gi1Q306K pull downs from ECs after flow, which was confirmed by Western blotting (Figure 2D). We next depleted the LPHNs in HUVECs. KD of LPHN2, but not LPHN1 or LPHN3, completely blocked endothelial alignment in flow and FSS-induced rapid responses (Figure 2E-H). Rescue by siRNA-resistant constructs revealed that all latrophilin isoforms rescue flow responses (Figure 2I and Extended data Fig S6), thus, are functionally equivalent. LPHN2 knockdown also blocked Gi2 activation by flow (Figure 2J & 2K). Thus, latrophilins are the flow-responsive GPCRs that regulate multiple PECAM1-dependent responses.

### Latrophilins in vascular development in zebrafish

To test the involvement of latrophilins in the vasculature *in vivo*, we examined zebrafish that express KDR-mCherry in ECs to mark these cells^23^. We used CRISPR-Cas9 with single guide (sg) RNAs that target all LPHN isoforms (1,2a, 2b and 3). Our designed sgRNAs showed >50% cleavage in T7 endonuclease assays as well as reduced mRNA levels due to nonsense-mediated decay (Extended data fig 7a and 7b). Embryos were stained for ZO-1 to mark cell boundaries. Combined KO of all of the LPHN isoforms inhibited alignment of ECs in the dorsal aorta (Figure 3A and 3B). Single knockouts revealed that LPHN2(a+b), but not LPHN1 or 3, similarly reduced EC alignment in the dorsal aorta (Figure 3C). To address whether this effect is due to defective flow sensing, we blocked blood flow by injecting single cell embryos with morpholinos against cardiac troponin T (TNNT or “silent heart”)^24^ at a dose that completely prevented cardiac contractility. TNNT morpholinos similarly blocked EC alignment; LPHN2 KO had no further effect, consistent with a role for LPHN2 in flow sensing (Figure 3D and 3E).

**Figure 3.**
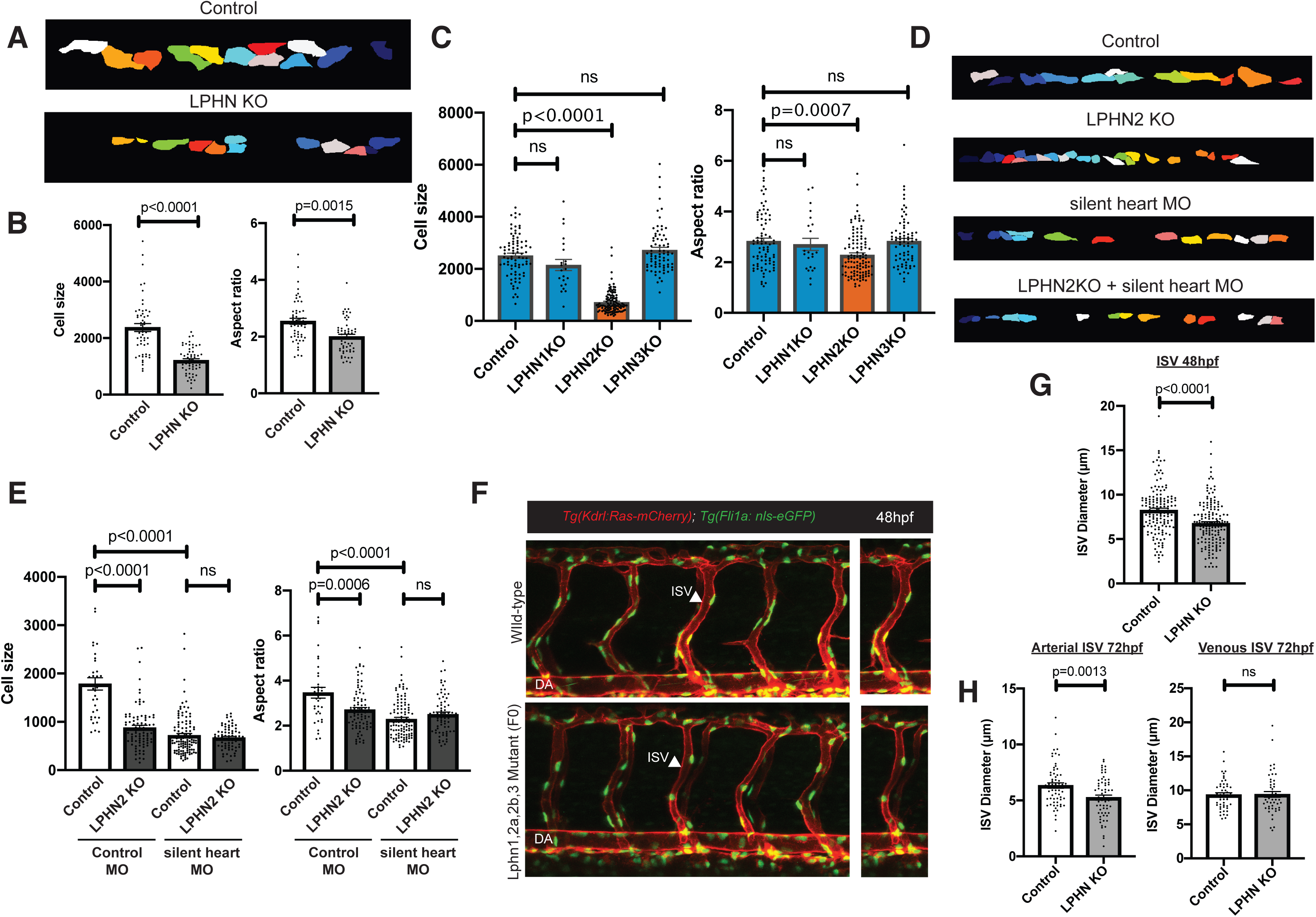
Latrophilin regulates endothelial flow responses *in vivo*. (**A**) Traces of endothelial shapes in the dorsal aorta at 48hpf from control zebrafish and with KO of all 4 LPHN isoforms. Data are representative of 6 fish. (**B**) Quantification of results from (A). Statistical analysis by two-tailed Student’s t-test. (**C**) Cell size and aspect ratio after knockout of individual latrophilins. LPHN2 KO includes 2a+b. Statistics analyzed by one-way ANOVA with Tukey’s multiple comparisons test. (**D**) Traces of endothelial cells from the dorsal aorta at 48hpf from embryos with or without LPHN2 KO and silent heart morpholinos (MO). Data are representative of 6 fish for each. (**E**) Quantification of (D). Statistical analysis by one-way ANOVA with Tukey’s multiple comparisons test. (**F**) Images of ISVs from control and LPHN KO embryos at 48 hpf. (**G**) Quantification of diameters from all ISV in (F). Comparisons were done by two-tailed Student’s t-test. (**H**) Quantification of diameters of arterial and venous ISVs at 72hpf of zebrafish. Data were obtained from 6 fish for each dataset. Data analysis was done by two-tailed Student’s t-test.

Artery lumen diameters are also determined by EC shear stress sensing^25^. At 48hpf, the diameters of intersegmental vessels (ISV) were reduced by both blockade of blood flow and latrophilin2 KO, with no further effect of combined inhibition (Figure 3F and 3G). When examined at 72hpf, when intersegmental arteries and veins can be distinguished, reduced diameter was only observed in the arteries (Figure 3H). Together, these data show that LPHN2 mediates EC flow sensing *in vivo*.

### Latrophilin-2 in FSS activation of Notch and Smads

FSS acting on ECs also activates Notch1 signaling at cell-cell contacts^26-28^. KD of LPHN2, but not LPHN1 or LPHN3, completely blocked the production of the Notch intracellular domain (ICD) in response to FSS (Figure 4A and 4B). Notch activation by its ligand, Dll4, was unaffected (Extended Data fig 8a&8b). Surprisingly, knockdown of Gi and Gq/11 had no effect (Figure 4C). Thus, LPHN2 is specifically required for Notch activation by flow independently of G*α* protein functions.

**Figure 4.**
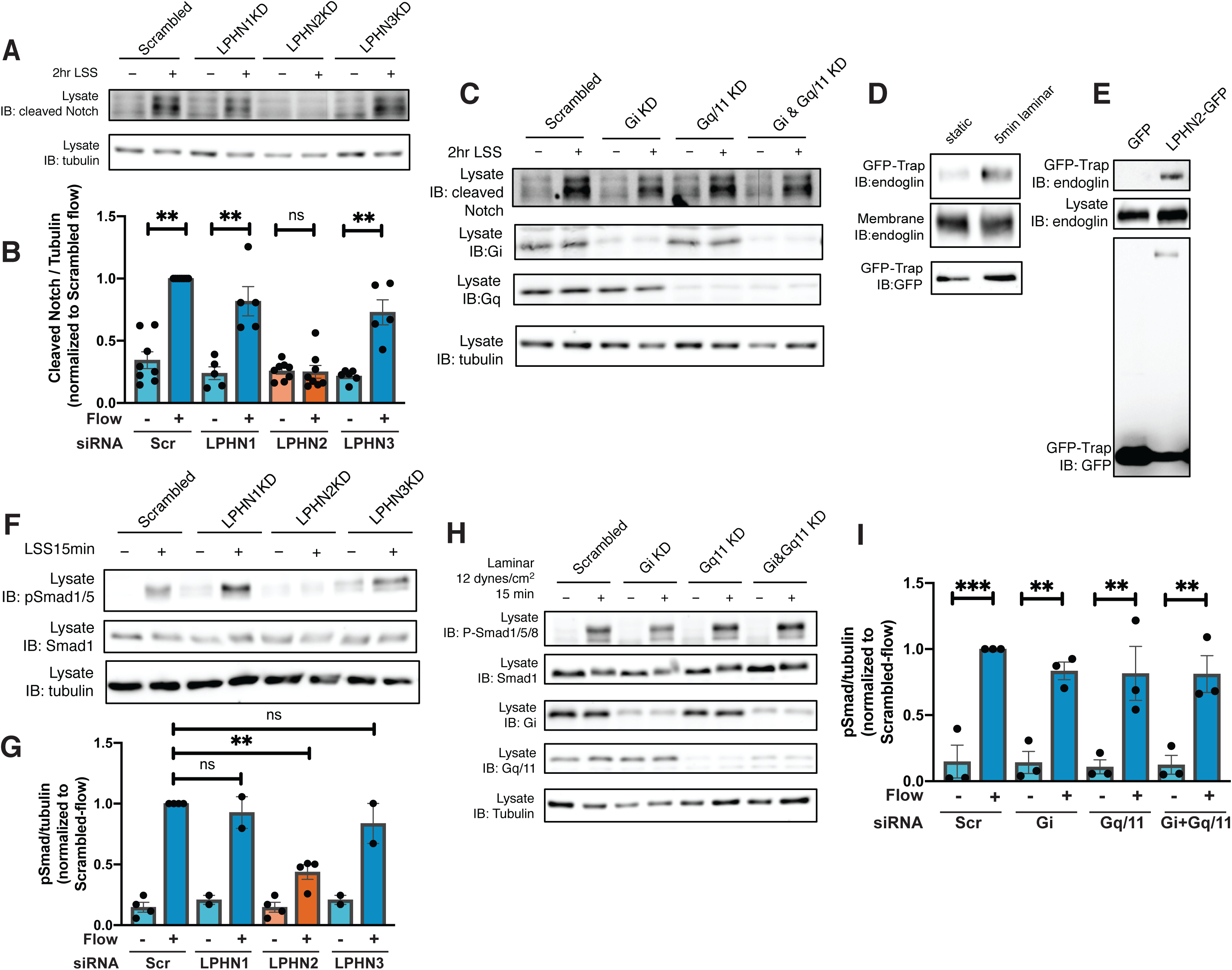
Flow-induced Notch and Smad1/5 signaling. (**A**) Western blot of cleaved Notch ICD in HUVECS with and without FSS, after KD of the indicated latrophilin isoforms. N=5. Results quantified in (**B**). N=6-8, **: p<0.001, one-way ANOVA with Tukey’s multiple comparisons test. (**C**) Western blot for cleaved Notch ICD with and without FSS after KD of the indicated G proteins. N=3. (**D**) Western blot for endoglin in Gi1Q306K pulldowns from cells with and without FSS. N=3 (**E**) HUVECs expression GFP or GFP-LPHN2 were immunoprecipitated with GFP-TRAP beads and analyzed by Western blotting for the indicated proteins. N=2. (**F**) HUVECs after KD of the indicated LPHN isoforms were subject to FSS and analyzed by Western blotting for pSmad1/5. N=4 for LPHN2, n=2 for LPHN1 and 3. Results quantified in (**G**). **: p<0.01, two-tailed Student’s t-test. (**H**) HUVECs after KD of the indicated G proteins were subject to FSS and Smad1/5 activation assayed as in F. Results quantified in (**I**). **: p<0.01, ***: p<0.001, one-way ANOVA with Tukey’s multiple comparisons test.

FSS also activates Smad1/5, which requires the BMP receptors Alk1 and endoglin, and occurs at cell-cell contacts^29^. Proteomic hits in the Gi1Q306K pull downs also contained endoglin, and Western blotting confirmed its FSS-depending association with Gi1306K and with LPHN2 complex (Figure 4D&4E). LPHN2 KD blocked activation of Smad1/5 by FSS (Figure 4F&4G) but had no effect on Smad1/5 activation by its soluble ligand BMP9 (Fig 4H&4I). KD of the G*α* proteins also had no effect on FSS activation of Smad1/5 (Extended data fig 7c&7d). By contrast, flow activation of Smad2/3 and induction of Klf2 did not require LPHN2 (Extended Data fig 9a-c).

## Discussion

These data identify the adhesion type GPCRs, latrophilins, as central organizers of EC flow mechanotransduction at cell-cell contacts. LPHNs mediate activation of the PECAM pathway through a G protein-dependent mechanism. Efficient inhibition required KD of both Gi2 and Gq/11, suggesting parallel or redundant effector pathways downstream of these G proteins. Curiously, previous studies identified either Gi or Gq/11 as essential for flow signaling to the same pathways, including PECAM, Akt, SFKs and eNOS^7,9,11^. It seems plausible that, depending on the EC type or experimental conditions, one or the other may predominate, concealing this functional redundancy. Understanding this GPCR-G protein signaling network is thus an important direction for future work. The novel pull downs assays and GPCR-G protein affinity purification protocol developed here may facilitate those efforts.

In contrast to PECAM, the role of LPHN in activation of Notch and Smad1/5 signaling is independent of Gi2 and Gq/11. For these pathways, the role of LPHNs appears to be to confer flow-sensitivity to these pathways. How LPHNs mediate this noncanonical, G protein-independent mechanisms is an open question.

KO of LPHN2 isoforms in zebrafish resulted in vascular defects consistent with FSS mechanotransduction. LPHNs have been hypothesized to mediate mechanotransduction in neurons^30^, though mechanistic insight is lacking. It is notable that, while LPHNs have been studied almost exclusively in the brain^31^, KO of LPHN2 in mice was embryonic lethal due to a non-neuronal function, consistent with an essential role in vascular development. However, single cell RNAseq studies^32^ detected other LPHN isoforms in non-neuronal tissues. Thus, LPHN1 or 3 may also contribute to mechanotransduction in different cell types or ECs in different tissues. These findings open the door to elucidation of their role in vascular development and disease, and in uncovering molecular mechanisms for converting physical force into biochemical signaling.

## Acknowledgements

This work was supported by NIH grant RO1 HL75092 to MAS, NIHLBI 5R01HL130246-05 to SN, and the Uehara memorial foundation postdoctoral fellowship and JSPS Overseas Research Fellowships (2014, receipt number: 805) to KT. We thank Hamit Harun Dag (Istanbul University) for help synthesizing SMA polymer; Shahid Mansuri (Yale University), Jeremy Herrera (University of Manchester) and David Knight (University of Manchester) for mass spectrometry experiments; Didier Trono (École polytechnique fédérale de Lausanne) for lentiviral packaging plasmids pMD2.G (Addgene plasmid # 12259; http://n2t.net/addgene:12259; RRID:Addgene_12259) and psPAX2 (Addgene plasmid # 12260; http://n2t.net/addgene:12260; RRID:Addgene_12260); Bryan Roth (University of North Carolina) for pcDNA5/FRT-HA-hM3D(Gq) (Addgene plasmid # 45547; http://n2t.net/addgene:45547; RRID:Addgene_45547) and pcDNA5/FRT-HA-hM4D(Gi)(Addgene plasmid # 45548; http://n2t.net/addgene:45548; RRID:Addgene_45548); Aziz Moqrich (Institut de Biologie du Dévelopement de Marseille) for GST-GINIP plasmid. We are grateful to Peter Newman (Medical College of Wisconsin) for the kind gift of PECAM antibody and helpful discussions.

## Figure Legends

**Figure S1.**
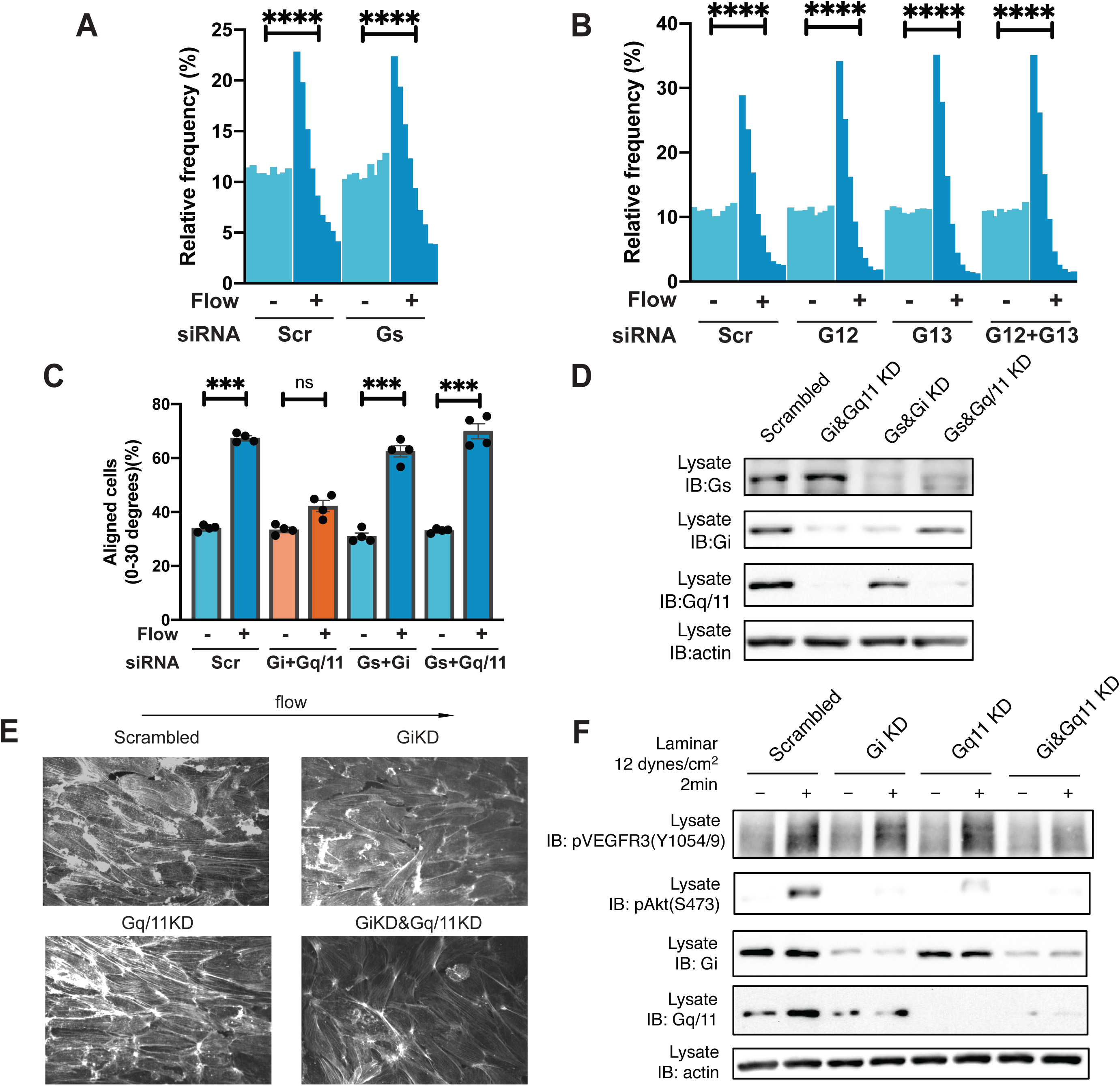
Gi and Gq/11 in flow responses. (**A**) HUVECs transfected with siRNAs targeting Gs were subjected to fluid shear stress (FSS) at 12 dynes/cm2 for 16 hours. Nuclear orientation was quantified to determine cell orientation to LSS. Alignment shown as a histogram as for Fig 1 and described in Methods. ****: p<0.0001, one-way ANOVA with Tukey’s multiple comparisons test. (**B**) HUVECs transfected with siRNAs for G12 or/and G13 were subjected to FSS and alignment determined as in A. ****: p<0.0001, one-way ANOVA with Tukey’s multiple comparisons test. (**C**) HUVECs transfected with siRNA targeting the indicated Gα proteins were subjected to FSS and alignment determined as before. ***: p<0.001, one-way ANOVA with Tukey’s multiple comparisons test. (**D**) Western blot for Gα proteins from (C). (**E**) Phalloidin staining of cells from Figure 1A. Cells were exposed to 12 dynes/cm^2^ laminar shear stress for 16 hours. (**F**) Activation of VEGFR3 and Akt in cells with knockdown of the indicated proteins with and without FSS.

**Figure S2.**
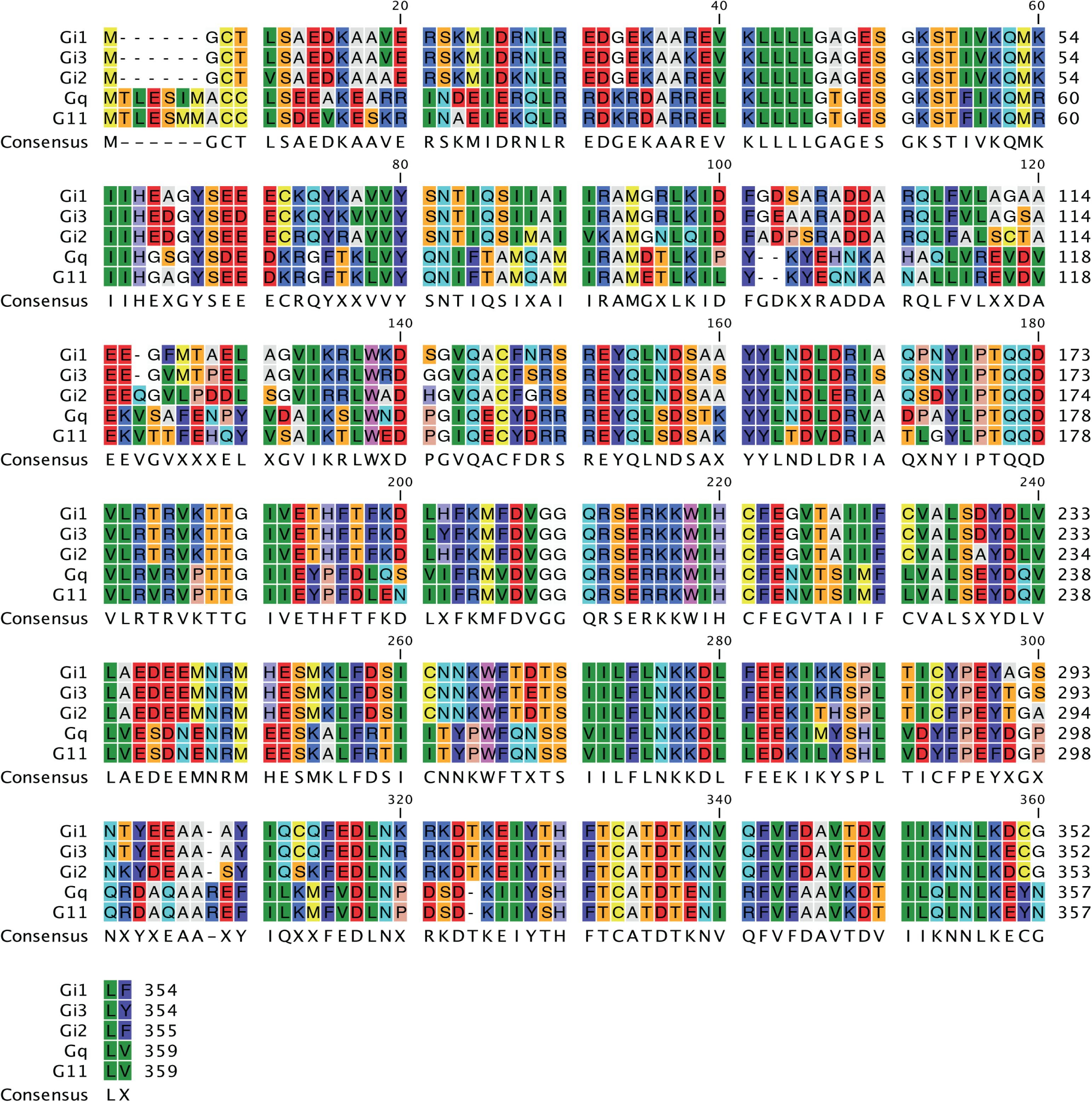
Amino acid sequence homology among G proteins. Complete sequence alignment of the indicated Gα proteins.

**Figure S3.**
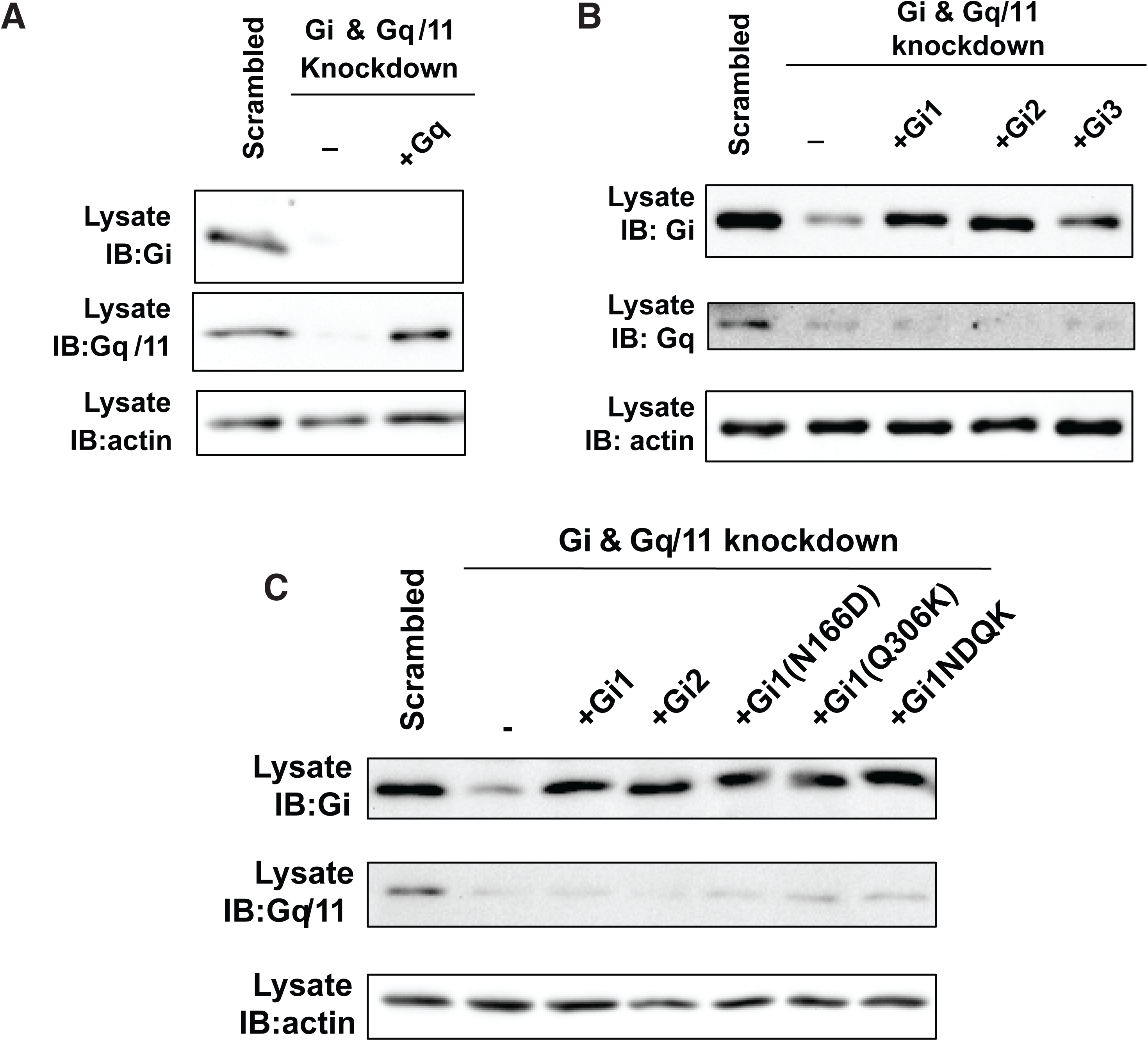
Confirmation of Gα KD and rescue for Figure 1. Expression level of Gq (**A**) and Gi (**B**) for the experiment in Figure 1D. (**C**) Expression level of Gi1, Gi2 and Gi1 mutants for the experiment in Figure 1F.

**Figure S4.**
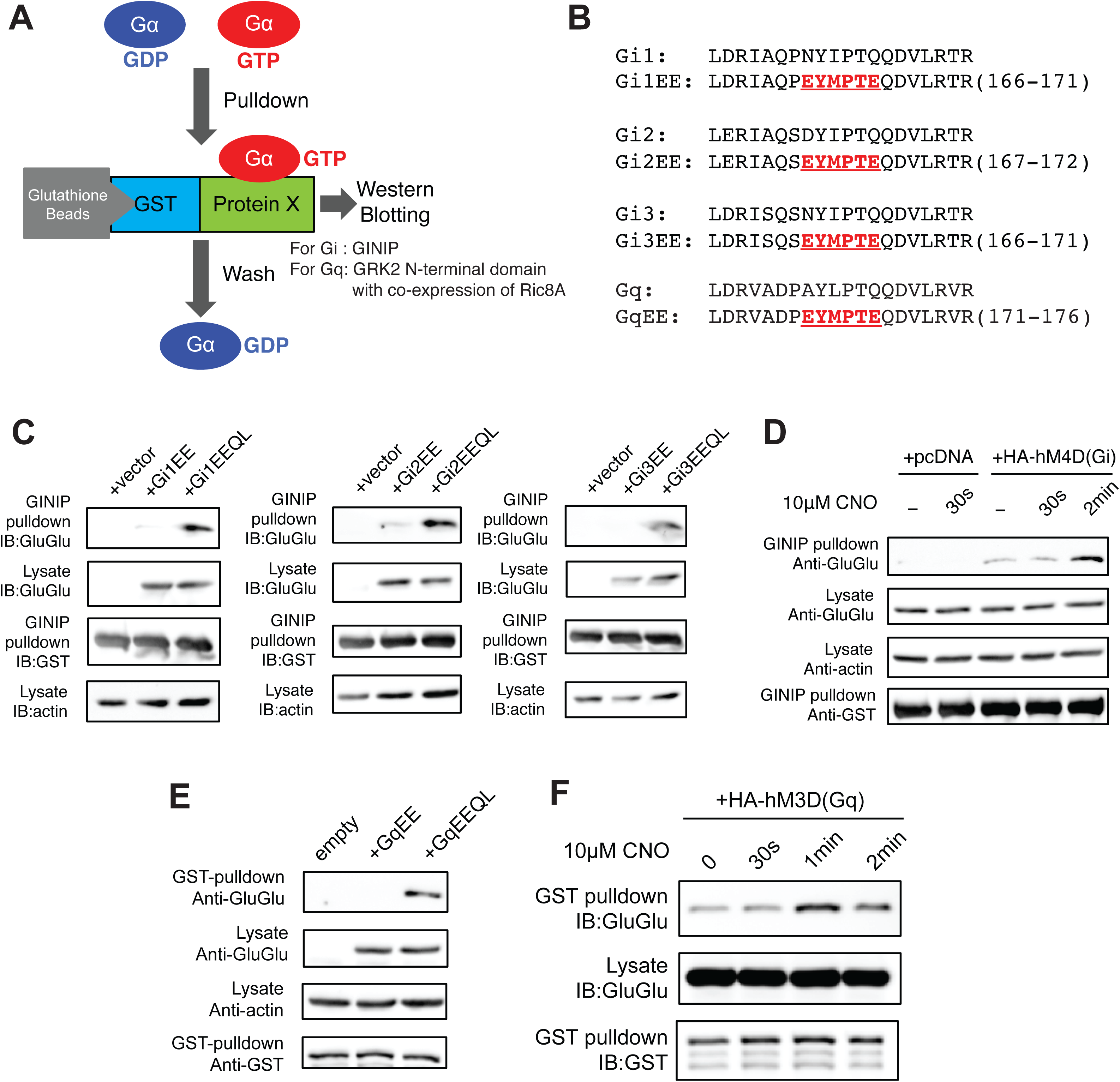
Pulldown assay for Gα activation. (**A**) Workflow for pull down assay for Gα activation. (**B**) Amino acid sequences showing the inserted GluGlu tag. (**C**) GINIP pulldown assay with WT and constitutively active Gi proteins. (**D**) GINIP pulldown assay for Gi activation by artificial DREADD M4D(Gi-specific) by its ligand 10*µ*M CNO. (**E**) GRK2N pulldown assay with constitutively active Gq protein. (F) GRK2 pulldown assay with artificial DREADD M3D(Gq-specific) after addition of 10*µ*M CNO.

**Figure S5.**
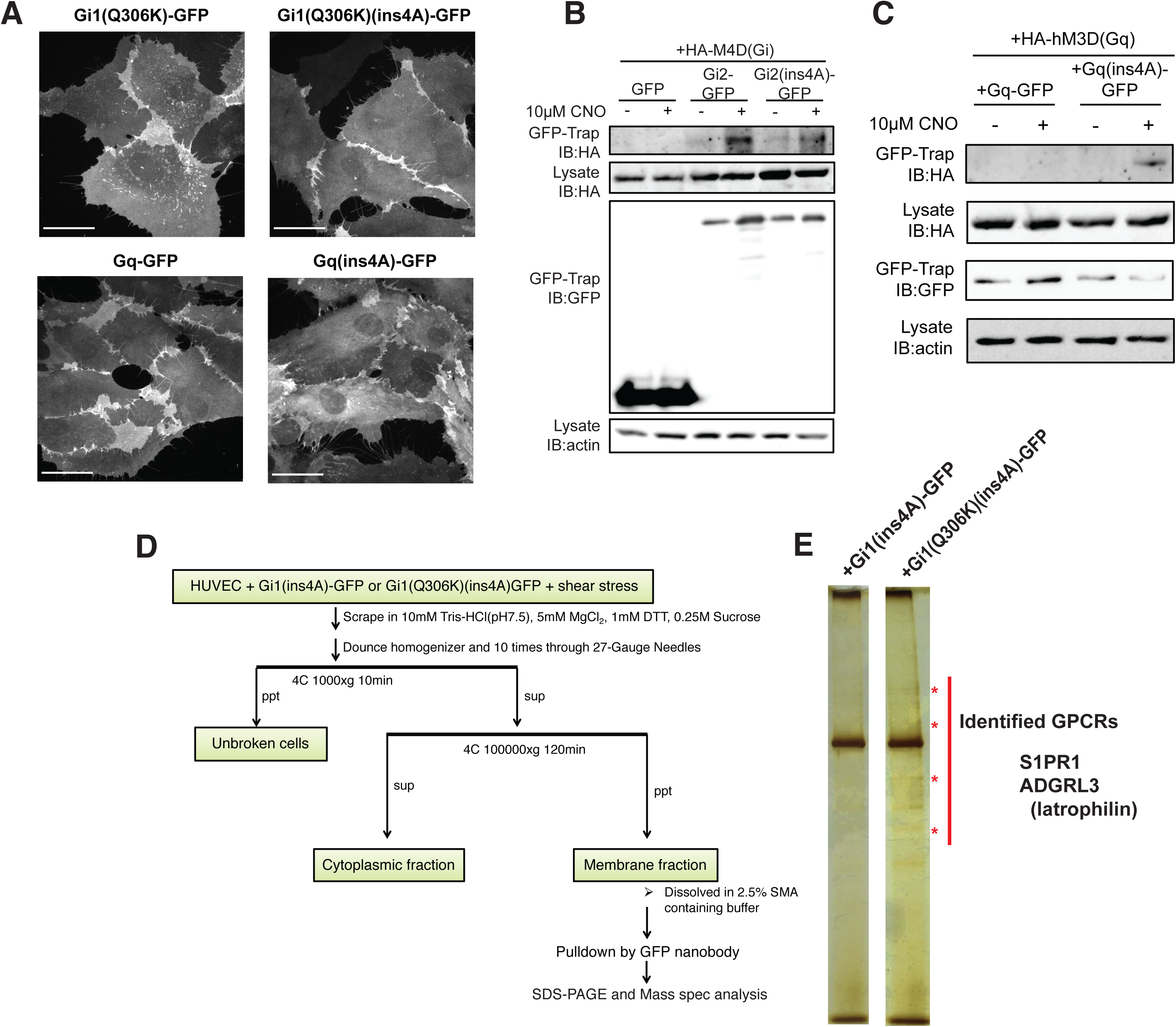
Proteomic identification of latrophilin. (**A**) HUVECs were infected with lentivirus expressing the indicated GFP-labeled G proteins and images taken by 60x objective on PerkinElmer spinning disk confocal microscope. Scale bar: 30*µ*m. (**B**) HEK293 cells expressing Gi2(ins4A)-GFP and HA-tagged M4D(Gi) were treated with or without 10 *µ*M CNO. Purified membrane fractions were solubilized by SMA polymer and immunoprecipitated with GFP-TRAP nanobody. IPs were analyzed by Western blotting for the indicated proteins. (**C**) HEK293 cells expressing Gq-GFP/Gq(ins4A)-GFP protein and HA-M3D(Gq) were treated with or without 10 *µ*M CNO. Membranes were isolated, extracted and analyzed as in B. (**D**) Workflow for proteomic identification of GPCRs. (**E**) Silver staining of isolated Gα(ins4A)-GFP membrane fractions from HUVECs with Gi1(ins4A)-GFP (negative control) or Gi1(Q306K)(ins4A)-GFP (samples). Red asterisks indicate the specific protein bands in Gi1(Q306K).

**Figure S6.**
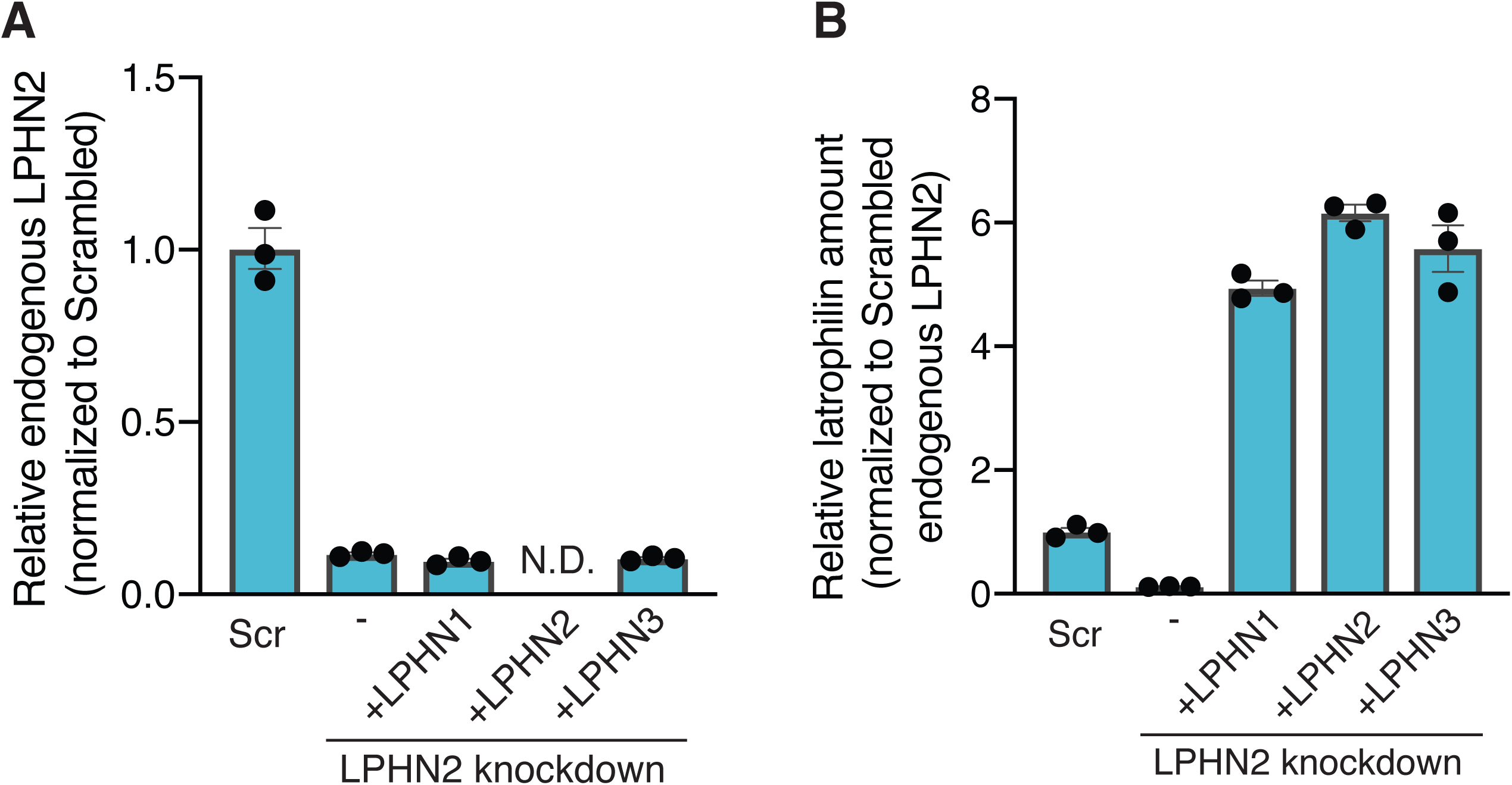
Confirmation of LPHN2 KD and rescue for Figure 2. mRNA expression level of endogenous LPHN2 (**A**) and LPHN rescue constructs (**B**) for the experiment in Figure 2I.

**Figure S7.**
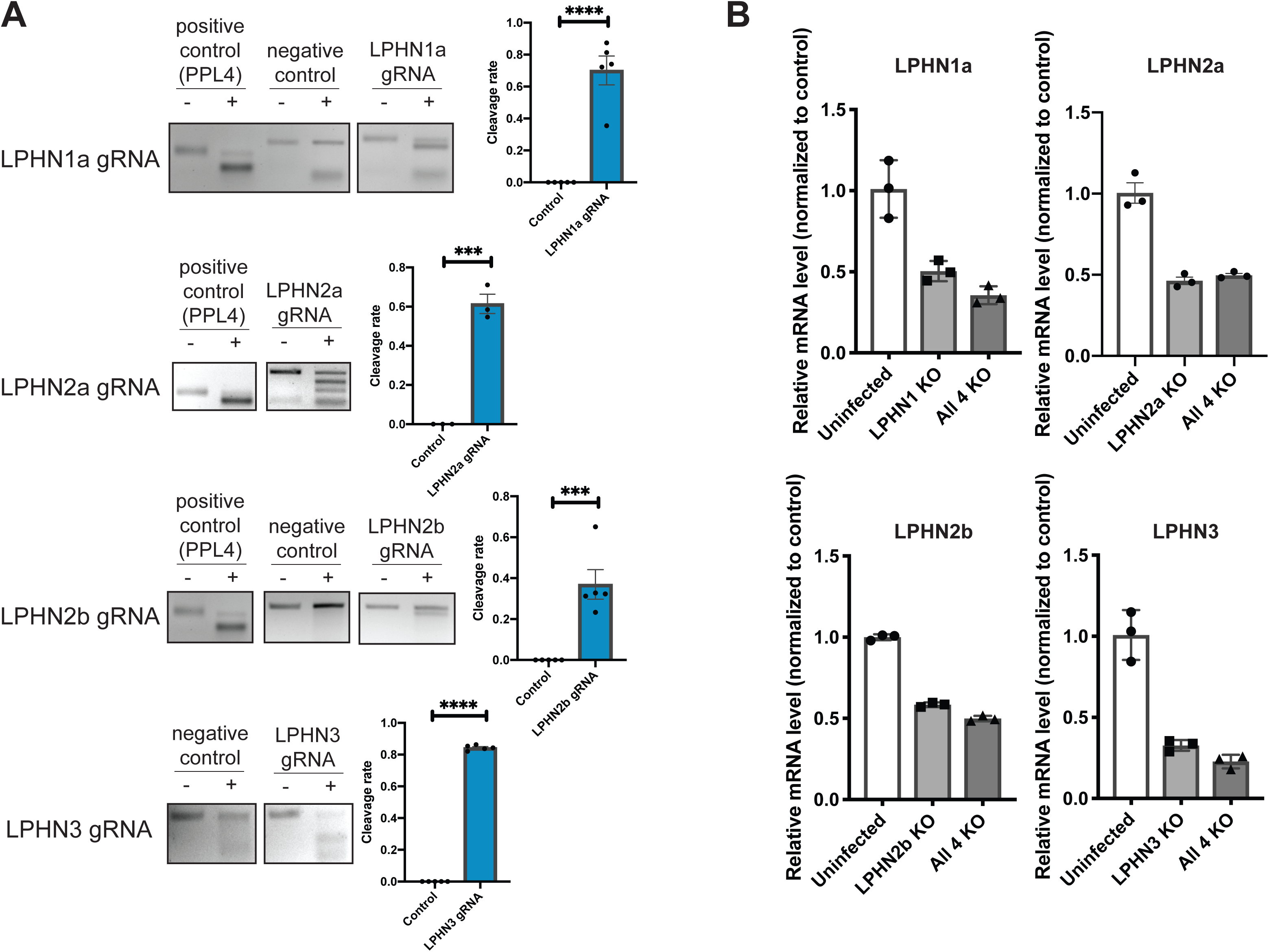
Validation of latrophilin knockout zebrafish. (**A**) T7 endonuclease assay for sgRNA targeting latrophilin isoforms. The positive control is the sample from zebrafish injected with sgRNA for ppl4. Negative control is zebrafish without injection. N=3-5 fish. ***: p<0.001, ****: p<0.0001, two-tailed Student’s t-test. (**B**) RT-PCR for indicated latrophilin isoforms at 48 hours after injection of sgRNA and mRNA encoding Cas9 endonuclease. Degradation of all isoforms due to nonsense mediated decay was observed.

**Figure S8.**
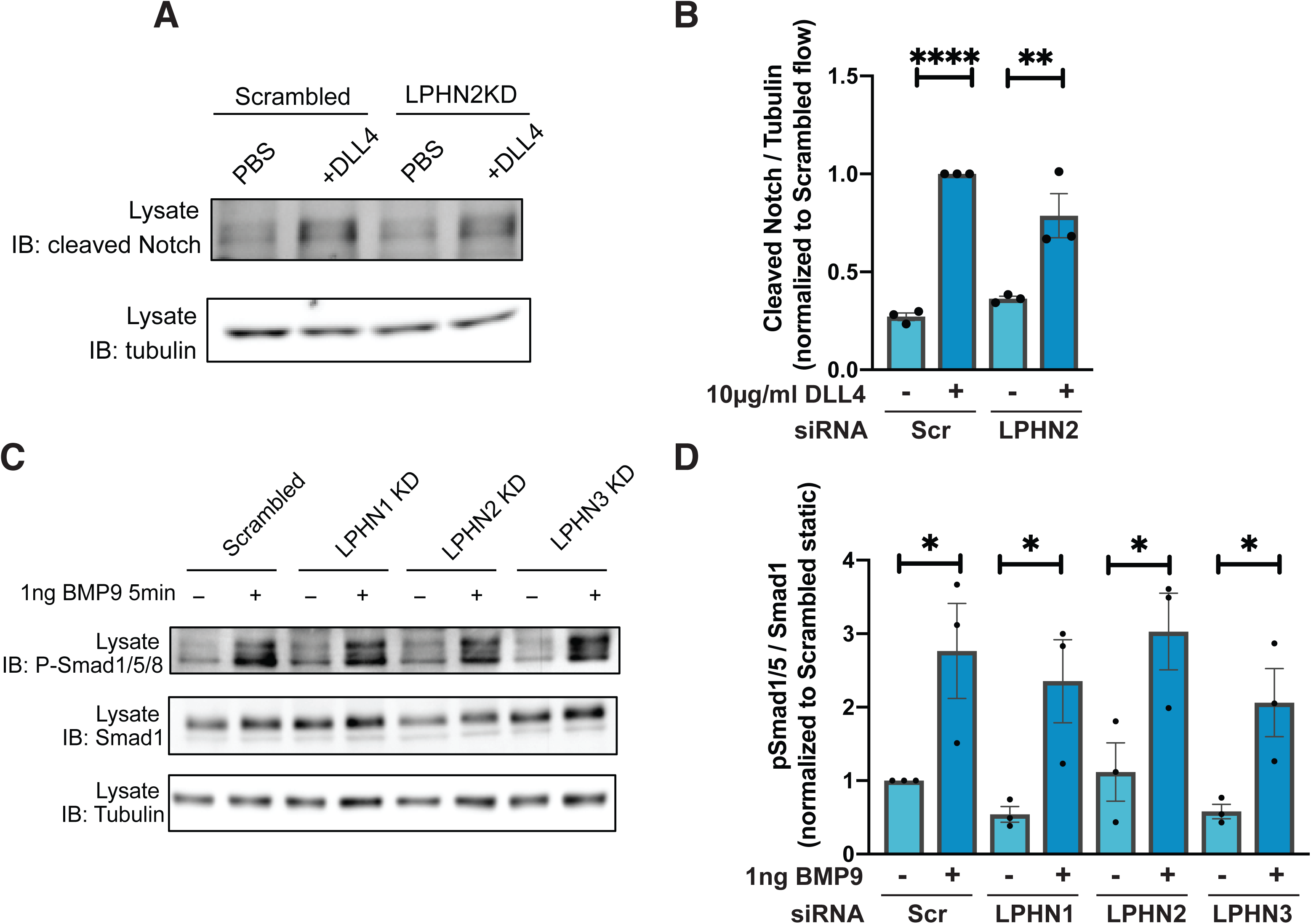
Ligand-induced activation of Notch signaling and Smad1/5. (**A**) HUVECs with or without LPHN2 KD were plated on dishes coated with DLL4 and Notch activation assayed by Western blotting. N=3. Results quantified in (**B**). **: p=0.0035, ****: p<0.0001, one-way ANOVA with Tukey’s multiple comparisons test. (**C**) HUVECs with or without KD of the indicated LPHNs were treated with 1ng BMP9 for 5 min and Smad1/5 activation assayed by Western blotting. N=3. Results quantified in (**D**). *: p<0.05, one-way ANOVA with Tukey’s multiple comparisons test.

**Figure S9.**
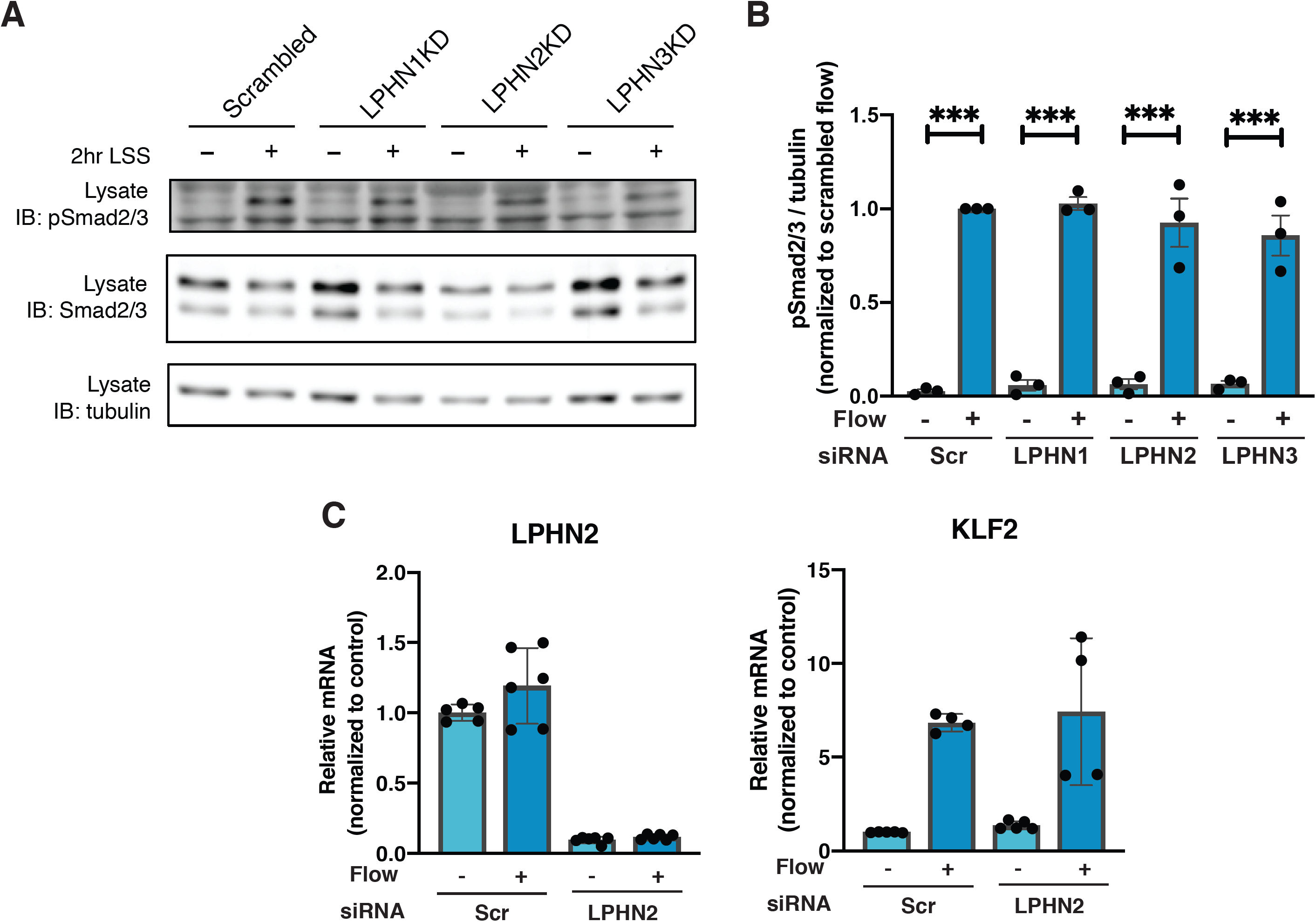
Latrophilin-independent FSS pathways. (**A**) HUVECs after KD of the indicated LPHNs were subjected to FSS and Smad2/3 activation assayed. N=3. Results quantified in (**B**). (**C**) ECs after KD of the indicate LPHNs were subjected to FSS for 16 hours and mRNAs were collected. Gene expression measured by RT-PCR. N=3.

## Experimental Procedures

### Antibodies

Primary antibodies used in this study were:

Anti-Phospho-Src (Tyr 416) (Cell signaling) (6943); Anti-Src (Cell signaling) (2109); Anti-Gi (NewEast Biosciences) (26003); Anti-Gq (NewEast Biosciences) (26060) and (BD Bioscience) (612705); Anti-actin (Santa Cruz) (sc-8432); Anti-Gs (NewEast Biosciences) (26006); Anti-Phospho-VEGFR3 (Tyr 1054/Tyr 1059) (Invitrogen) (44-1047G); Anti-Phospho-Akt (Ser 473) (Cell Signaling) (9271); Anti-GluGlu (BioLegend) (MMS-115P); Anti-GST (Cell signaling) (2625); Anti-Tubulin (Invitrogen) (62204); Anti-GFP (Santa Cruz) (sc-9996) and (Abcam) (ab13970); Anti-HA (BioLegend) (901501); Anti-VE-Cadherin (Santa Cruz) (sc-6458); Anti-RFP (Antibodies-Online) (ABIN129578); Anti-ZO-1 (Invitrogen) (61-7300); Anti-cleaved Notch (Cell Signaling) (4147); Anti-Endoglin (R&D Systems) (AF1097); Anti-Phospho-Smad1/5 (Cell signaling) (13820); Anti-Smad1 (Cell signaling) (9743); Anti-Phospho-Smad2/3 (Cell signaling) (8828); Anti-Smad2/3 (Cell signaling) (8685); Anti-PECAM-1 was kind gift from Peter Newman, Milwaukee Blood Center.

### Cell Culture

Primary HUVECs were obtained from the Yale Vascular Biology and Therapeutics core facility. Each batch is composed of cells pooled from three donors. Cells were cultured in M199 (Gibco: 11150-059) supplemented with 20% FBS, 1x Penicillin-Streptomycin (Gibco: 15140-122), 60 *µ*g/ml heparin (Sigma: H3393), and endothelial growth cell supplement (hereafter, complete medium). HUVECs used for experiments were between passages 3 and 6.

### Shear stress

HUVECs were seeded on tissue culture-treated plastic slides coated with 10 *µ*g/ml fibronectin for 1 hour at 37°C and grown to confluence. For short-term experiments, cells were starved overnight in M199 medium with 2% FBS and 1:10 of ECGS or for 30 minutes in M199 medium containing 0.2 % BSA. These slides were set in parallel flow chambers and shear stress applied as described^33^.

### Image analysis

Cell orientation was calculated by taking the masks of the cell nuclei determined by Hoechst images, fitting them as an ellipse, and determining the angle between flow direction and the major axis of the ellipse. Analyzed results were visualized as histograms showing the percent of cells within each 10° of the direction of flow or as quantification of aligned cells with nuclei whose major axis were within 0-30 degrees to flow direction.

### Data display

All data displayed were means ± SEM except n=2, in which case they were means ± range.

### Lentiviral transduction

Lenti-X 293T cells (Clontech, 632180) were cultured for at least 24 hours in DMEM supplemented with 10% FBS and lacking antibiotics, then transfected with lentiviral plasmids encoding the gene of interest and packaging plasmids (Addgene: 12259 and 12260) using Lipofectamine 2000 (Thermo Fisher Scientific: 11668-019) following the manufacturer’s protocols with Opti-MEM medium. Conditioned media from these cultures were collected 48 hours later, sterilized through 0.22*µ*m filters and added to HUVECs together with 8*µ*g/ml of polybrene (Sigma: 107689). After 24 hours, cells were switched to complete medium for 48 hours.

### siRNA transfection

HUVECs were cultured in EGM™-2 Endothelial Cell Growth Medium-2 BulletKit™ (Lonza: CC-3156 and CC-4176) for 24 hours before transfecting with RNAiMax (Thermo Fisher Scientific: 13778-150) with 20nM siRNA in Opti-MEM (Gibco: 31985-070) according to the manufacturer’s instructions. After 6 hours, cells were switched to EGM-2 medium and used for experiments 2-3 days later. Gα protein siRNAs were as described previously^34-36^. For latrophilin knockdown, ON-TARGET plus Smartpool siRNAs from Dharmacon against human LPHN1 (L-005650-00-0005), LPHN2 (L-005651-00-0005), LPHN3 (L-005652-00-0005) were used.

### Western Blotting

HUVECs were washed with PBS and extracted in Laemmli sample buffer. Samples were separated by SDS-PAGE and transferred onto nitrocellulose membranes. Membranes were blocked with 5% milk in TBS-T and probed with primary antibodies at 4°C for overnight. The targeting proteins were visualized by HRP-conjugated secondary antibodies and subsequent HRP-luminol reaction.

### Immunofluorescence

HUVECs were washed with PBS, fixed for 10 minutes with 3.7% PFA in PBS. Following fixation, cells were permeabilized with 0.5% Triton X-100 in PBS for 10 minutes and then incubated with 3% BSA in PBS for 30 minutes for blocking. Cells were washed with PBS after blocking and were incubated Alexa488-Phalloidin and Hoechst for 1 hour, then washed 4 times with PBS and mounted. Images were captured with 20x or 60x objective on a PerkinElmer spinning disk confocal microscope. Cell alignment was determined as described previously^37^.

### Reverse transcription and quantitative PCR

RNA was isolated from HUVECs using RNeasy kit according to the manufacturer’s instructions and quantified using a nanodrop spectrophotometer. Following cDNA synthesis using Bio-Rad iScript kit, RT-PCR was performed as follows. Each PCR reaction contains 42 two-step amplification cycles consisting of: (a) denaturation at 95°C for 30s, and (b) annealing and extension at 60°C for 30s. The amplification curve was collected, and the relative transcript level of the target mRNA in each sample was calculated by normalization of Ct values to the reference mRNA (GAPDH). Primer sequences used for RT-PCR were as shown in Table 1.

**Table 1:**
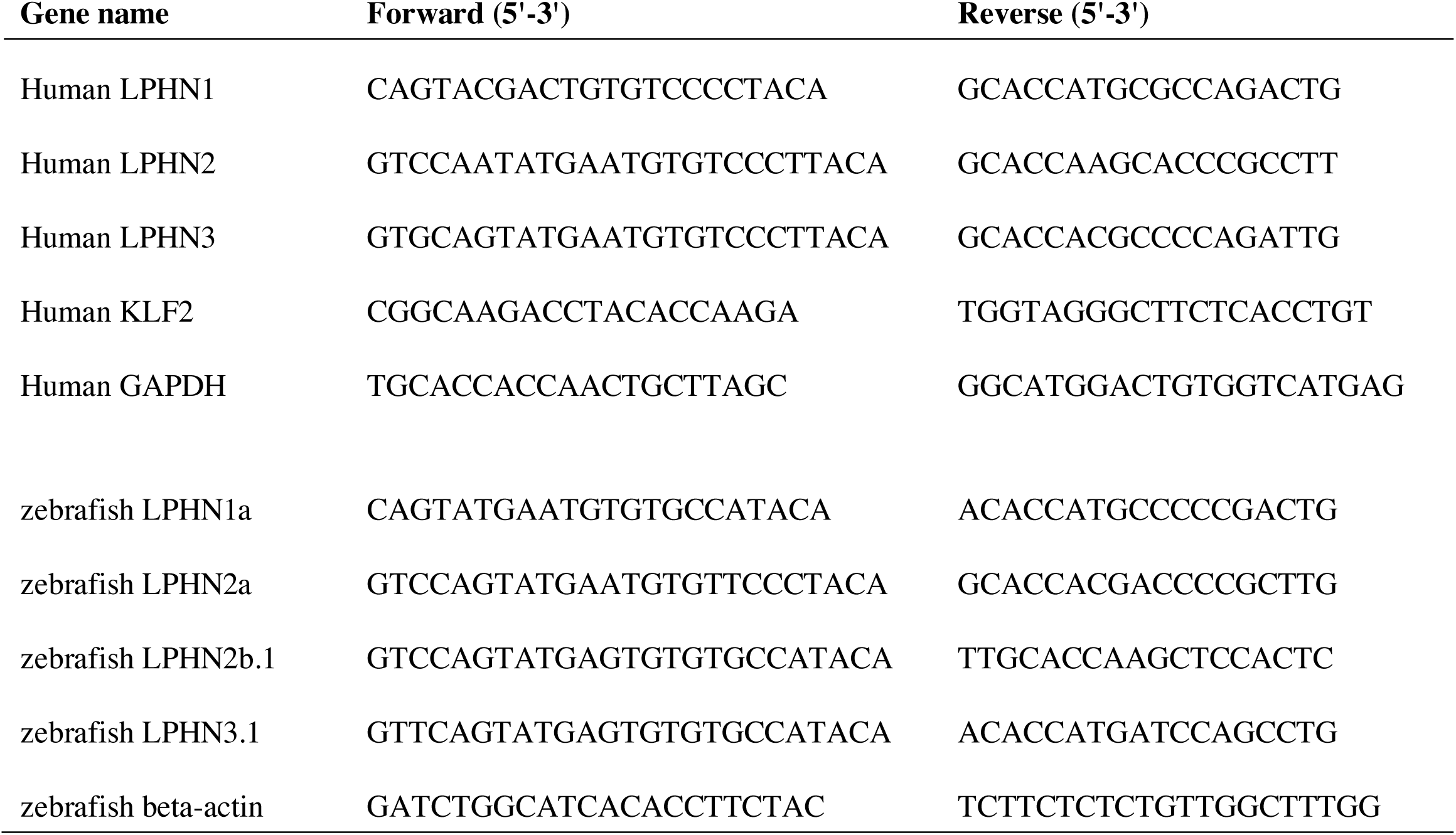
Primer sequences used for quantitative PCR in this study.

### G protein pull-down assay

BL21 *E.coli* cells were transformed with constructs expressing GST-tagged GINIP protein or GST-tagged GRK2 N-terminal domain. Cells were incubated in terrific broth (http://cshprotocols.cshlp.org/content/2015/9/pdb.rec085894.full?rss=1) and protein expression induced by addition of 0.5*µ*g/ml IPTG. Cells were collected after 8 hours, lysed and GST-proteins collected on Glutathione-conjugated beads. Beads were washed 4 times, eluted with 50mM glutathione and proteins desalted on a gel filtration column. Aliquots of 20 *µ*g aliquots were stored frozen and thawed shortly before use.

For pulldown assays, HUVECs with GluGlu-tagged Gα mutant were lysed in 10mM Tris-HCl pH7.5, 150mM NaCl, 1% Triton X-100 and 5mM DTT and lysates were incubated for 10 minutes at 4°C with gentle agitation with glutathione beads pre-conjugated with 20*µ*g of GST-GINIP or GST-GRK2N. Beads were washed three times with cell lysis buffer, solubilized in SDS sample buffer, and analyzed by Western blotting.

### Zebrafish husbandry and handling

Zebrafish were housed and maintained at 28.5°C in accordance with standard methods approved by the Yale University Institutional Animal Care and Use Committee (#2017-11473)^38,39^.

### Generation of lphn/adgrl knockdown zebrafish using CRISPR/Cas9 ribonucleoproteins

Four zebrafish latrophilin paralogs were identified—*adgrl1a, adgrl2a, adgrl2b.1*, and *adgrl3.1*. Fused sgRNAs were generated targeting each paralog (see Table 2) by annealing locus-specific oligonucleotides to a common 5’ universal oligonucleotide and performing *in vitro* RNA transcription (AmpliScribe T7-Flash Kit, Lucigen, ASF3507)^40^. An injection mix of 50 ng/*µ*L each sgRNA, an equivalent concentration of Cas9 protein (TrueCut Cas9, Invitrogen, A36497), and either 1 *µ*g/*µ*L *silent heart/tnnt2a*^*24*^ or standard control morpholino (GeneTools) was prepared. 1 *µ*L of the mixture was injected into *Tg(kdrl:ras-mCherry, fli1a:nls-GFP)* zebrafish at the one cell stage (Parker Hannifin, Picospritzer III). CRISPR reagent efficacy was confirmed by T7 endonuclease assay (see Extended data Fig 6). Guides against adgrl2a and 2b.1 were combined to target all LPHN2 isoforms.

**Table 2:**
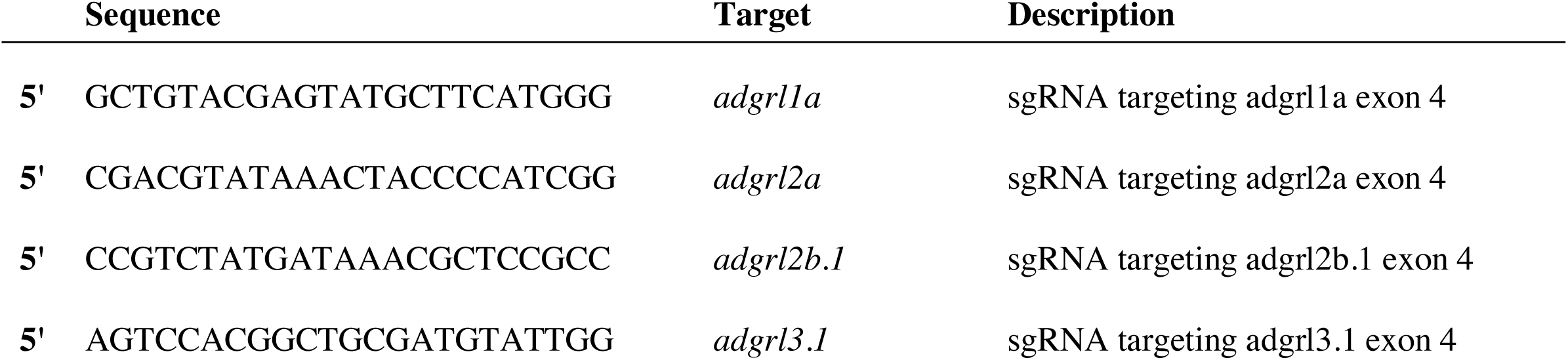
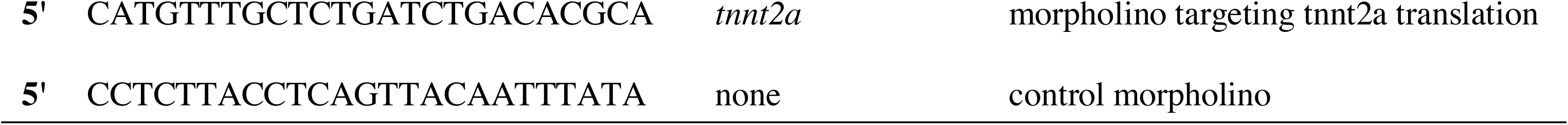
Sequences of sgRNAs and morpholinos used in this study.

### Immunostaining and imaging of knockdown zebrafish

CRISPR/Cas9 RNP-injected embryos were either fixed at 48 hpf in 4% paraformaldehyde (Santa Cruz Biotechnology, SC-253236) and immunostained for GFP (1:300 chicken anti-GFP, abcam, ab13970), mCherry (1:300 rabbit anti-RFP, Antibodies-Online, ABIN129578), and ZO-1 (1:200 mouse anti-ZO-1, Invitrogen, 61-7300) with species-appropriate secondary antibodies (1:400 Invitrogen anti-chicken/-rabbit/-mouse Alexa 488/546/647, A-11039/A-10040, A-31571) or imaged live. For fixed imaging, the larvae were processed as previously described^25^. For live imaging, larvae were anesthetized by immersion in 600 *µ*M tricaine methylsulfonate (Western Chemical, TRS5), and embedded in 1.5% low-melt agarose (BioRad, 1613106). Intersegmental vessel and dorsal aorta diameter were then calculated using ImageJ. All imaging was performed using a Leica SP8 confocal microscope.

